# Compensation of compromised PRC2 regulation by a miRNA ensures robustness of Arabidopsis leaf development

**DOI:** 10.1101/2023.10.11.561475

**Authors:** Aude Maugarny, Aurélie Vialette, Bernard Adroher, Nathalie Mathy-Franchet, François Roudier, Patrick Laufs

## Abstract

Robustness is pervasive throughout biological systems, enabling them to maintain persistent outputs despite perturbations in their components. Here, we reveal a novel mechanism contributing to leaf morphology robustness in the face of genetic perturbations. In Arabidopsis, leaf shape is established during early development through the quantitative action of the *CUP-SHAPED COTYLEDON2* (*CUC2*) gene that is negatively regulated by the co-expressed *MICRORNA164A* (*MIR164A*) gene. Compromised epigenetic regulation due to defective Polycomb Repressive Complex 2 (PRC2) function results in the transcriptional derepression of *CUC2* but has no impact on CUC2 protein dynamics or early morphogenesis. We solve this apparent paradox by showing that compromised PRC2 function simultaneously activates a compensatory mechanism involving another member of the *MIR164* gene family, the *MIR164B* gene. This mechanism dampens CUC2 protein levels, thereby compensating for compromised PRC2 function and canalizing early leaf morphogenesis. Furthermore, we show that this compensation mechanism is active under different environmental conditions. Our findings shed light on how the interplay between different types of transcriptional regulation can contribute to developmental robustness.

## Introduction

Biological systems are characterized by their robustness, which enables them to produce a constant output despite environmental or endogenous perturbations that affect the activity of individual components of the system (Félix and Barkoulas, 2015; Kitano, 2004). The structure of the biological system, for instance the presence of positive or negative feedback loops contributes to its robustness (Kitano, 2004). In the case of genetic networks, these regulatory interactions may operate through different levels of regulation such as transcriptional, post-transcriptional and post-translational regulations. For instance, miRNA negative regulation of gene expression at the post-transcriptional level has been suggested to play prominent roles in the robustness of the expression of target genes (Alberti and Cochella, 2017; Avital et al., 2018; Ebert and Sharp, 2012; Hornstein and Shomron, 2006; Posadas and Carthew, 2014). Transcriptional repression through epigenetic modifications of chromatin organisation by Polycomb Group (PcG) proteins, and in particular the Polycomb Repressive Complex 2 (PRC2,) has also been argued to be involved in genetic robustness (Elgart et al., 2015). Yet, how negative gene regulations provided by miRNAs and PRC2 may interact to contribute to robustness is far less understood.

PRC2 transcriptional and miRNA post-transcriptional gene regulations have been shown to be interconnected through different ways. A first mode of interaction is an antagonism between actors of the PRC2 and miRNA pathways resulting from their reciprocal inhibition (Moutinho and Esteller, 2017; de Nigris, 2016; Wang et al., 2015). Such double negative feedback loops can lead to bistable systems contributing to cell fate decision (Juan and Sartorelli, 2010). In plants, no miRNA regulating PcG-encoding genes have been identified so far, but miRNA genes are more frequently targeted by PRC2 regulation than protein coding genes (Lafos et al., 2011). Regulation of miRNA gene expression by PRC2 is particularly important for the fine tuning of plant progression from vegetative to reproductive phase (Picó et al., 2015; Xu et al., 2016). In a second type of interaction, PRC2 and miRNAs cooperate effect to provide additive repression at the transcriptional and posttranscriptional levels on a shared target gene. In glioblastoma multiform cells, about 20% of all direct PRC2-repressed genes are also repressed by miRNAs and in about half of these cases, the PRC2 and miRNA actions are coordinated as PRC2 indirectly promotes miRNA expression (Shivram et al., 2019). In plants, genes targeted by a miRNA tend to be also more frequently targeted by PRC2 than non-miRNA targets (Lafos et al., 2011), but examples of cooperation between PRC2 and miRNAs are still lacking. Here we reveal a novel mode of interaction between PRC2 and miRNA, in which a novel miRNA node compensates for compromised PRC2 function during Arabidopsis leaf development.

Leaves in plants are diverse in size and shape and this diversity is determined by genetic, developmental and environmental factors (Chitwood and Sinha, 2016). All leaves are initiated as small, finger-shaped structures and their complex shape is progressively set up through differential growth controlled by molecular and mechanical regulations. Leaf shaping is divided into two phases (Bar and Ori, 2014; Maugarny-Calès and Laufs, 2018). During the primary morphogenesis phase, the basic architecture is set up by the initiation and growth of teeth or leaflets along the leaf margin. During the secondary phase, this pattern can be rearranged, for instance by fusion between leaflets (Champagne et al., 2007), and the leaf expands. The Arabidopsis CUC2 transcription factor, like its homologues in other plant species, is a major regulator of the primary morphogenesis phase by having a dual role (Berger et al., 2009; Blein et al., 2008; Cheng et al., 2012; Nikovics et al., 2006; Zheng et al., 2019). First, through the interaction with auxin, CUC2 expression patterns the leaf margin to determine the number and position of the Arabidopsis teeth (Bilsborough et al., 2011; Kawamura et al., 2010). Next, CUC2 induces tooth outgrowth through a combination of cell autonomous repression of growth to form the teeth sinuses and a promotion of growth at distance at the tooth tip (Biot et al., 2016; Kierzkowski et al., 2019; Maugarny-Calès et al., 2019; Nikovics et al., 2006). Hence, leaf morphometry has shown that leaf dissection is a quantitative, morphological read-out of CUC2 protein levels in the teeth sinus (Maugarny-Calès et al., 2019). More precisely, leaf dissection can be observed on individual teeth and quantified as the tooth aspect ratio (TAR=tooth height/tooth width) or at the organ level as the leaf dissection index (LDI) to take into account repeated tooth formation along the leaf margin (Maugarny-Calès et al., 2019; Sicard et al., 2014) (Figure 1a).

**Figure 1:**
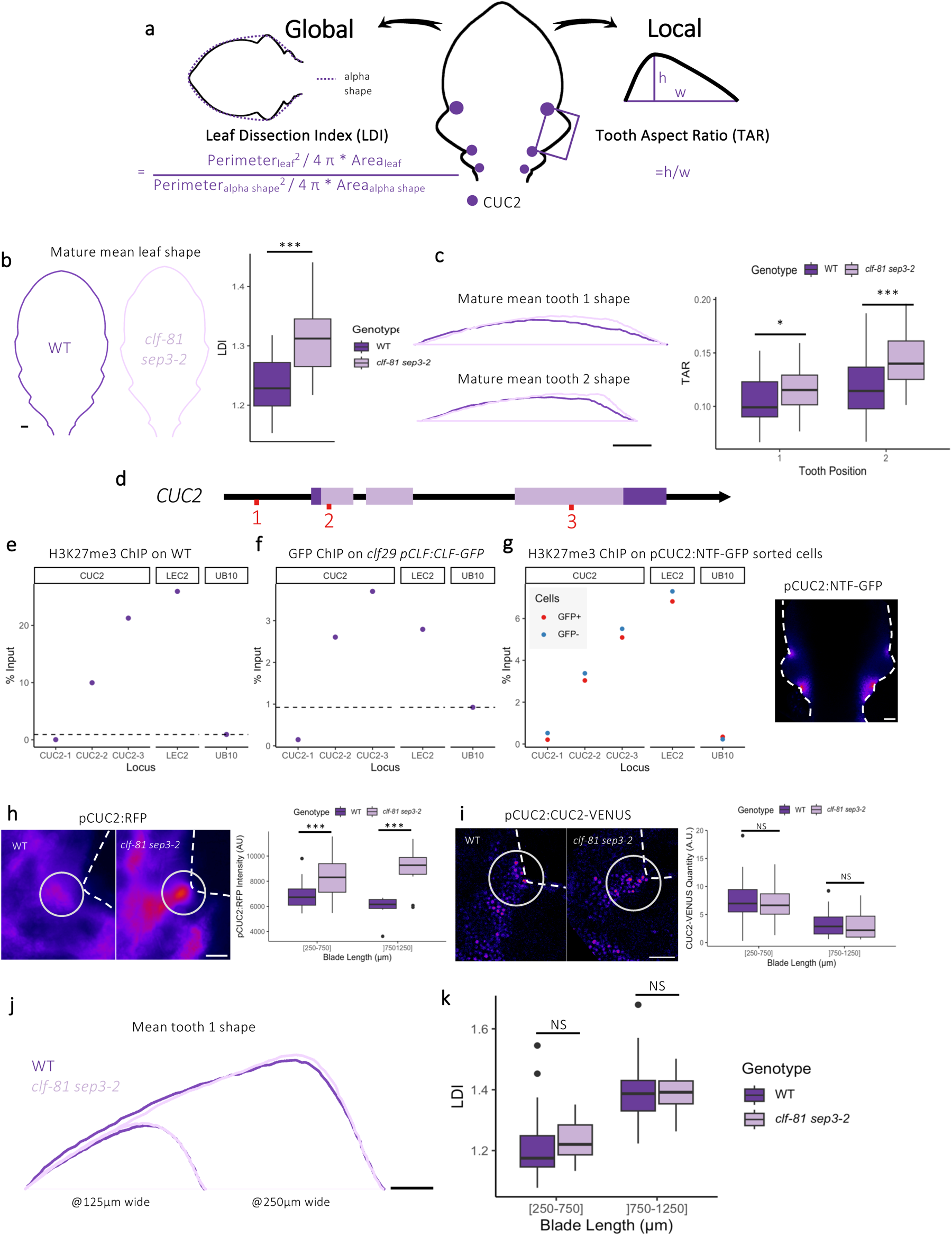
*CUC2* is derepressed in *clf-81 sep3-2* but dampened by an unknown mechanism. a) CUC2 expression and leaf morphometrics. *CUC2* expression patterns are shown by purple discs on the central leaf primordium. Tooth 1 in marked by a purple rectangle. Leaf dissection can be measured at the organ level as the Leaf Dissection Index (LDI) or at the tooth level as the Tooth Aspect Ratio (TAR). b) Mean leaf shape and LDI and c) Mean tooth shape and TAR of WT and *clf-81 sep3-2* mature leaves. *n* ≥ 20 in b) and *n* ≥ 36 in c). d) Schematic *CUC2* gene structure. Exons are represented by light purple boxes (translated regions) and dark purple boxes (untranslated regions), introns and promoter regions are represented by black lines. The amplified regions for ChIP-qPCRs in e), f) and g) are defined by the three sets of primers designated in green. e) H3K27me3 chromatin-immuno-precipitation followed by qPCR (ChIP-qPCR) on WT rosette centers. f) GFP ChIP-qPCR on *clf-29* pCLF:CLF-GFP rosette centers. The results of a single experiment are shown, the results of other biological replicates are available in Figure S2a-c. g) H3K27me3 ChIP-qPCR on pCUC2:NTF-GFP rosette centers GFP sorted nuclei. In g), the picture shows the expression pattern in a young leaf of the pCUC2:NTF-GFP reporter that was used to sort out GFP+ and GFP-cells. Dashed lines mark the leaf margin. The *LEC2* locus is known to be marked by H3K27me3 in this tissue (positive control), while the *UB10* locus acts as a negative control. h) pCUC2:RFP and i) pCUC2:CUC2-VENUS intensities in WT and *clf-81 sep3-2* distal sinus of tooth 1. h) and i) Images on the left are representative of the typical relative intensities. Dashed lines mark the leaf margin limit, the grey circles the quantification areas. Quantifications are plotted in arbitrary unit (A.U.) for two bins of blade length, *n* ≥ 8 for h) and *n* ≥ 6 for i). j) Mean shape of tooth 1 in WT and *clf-81 sep3-2* developing leaves. Small and large teeth are 125µm and 250 µm wide, respectively, which correspond to leaf primordia about 500µm and 1000µm long. k) LDI of WT and *clf-81 sep3-2* developing leaves. *n* ≥ 31. Statistical significance is tested by Student’s tests: NS for not significant, p-value is <0.05 for *, <0.01 for **, <0.005 for ***. Scale bars in g), h), i) and j) is 30 µm, in b) and c) 1 mm.

CUC2 activity is regulated at multiple levels: transcriptionally (Bilsborough et al., 2011; Galbiati et al., 2013; Nicolas et al., 2022), post transcriptionally by microRNA miR164 (Larue et al., 2009; Laufs et al., 2004; Nikovics et al., 2006) and via protein interactions (Barro-Trastoy et al., 2022; Gonçalves et al., 2015; Rubio-Somoza et al., 2014). In Arabidopsis, miR164 is encoded by three genes. *MIR164A* is co-expressed with *CUC2* at the leaf margin where their interaction controls CUC2 levels and leaf dissection (Maugarny-Calès et al., 2019; Nikovics et al., 2006). *MIR164C* plays a similar role in flowers where it limits the level of CUC2 and of its paralogue CUC1 to control floral organ patterning (Baker et al., 2005). Finally, *MIR164B* is more broadly expressed and the main contributor of miR164 at the whole seedling level, but its specific role is not known (Mallory et al., 2004). Here, we show that *CUC2* is also negatively regulated by PRC2, but that surprisingly no changes in CUC2 protein levels nor early leaf shape are observed in mutants with defective PRC2 function. We reconcile these conflicting observations by uncovering a compensatory mechanism involving the activation of *MIR164B* in leaves with compromised PRC2 function. The activation of *MIR164B* provides robustness to CUC2 protein expression and early leaf development despite variations in *CUC2* transcription. Furthermore, we show that this compensatory mechanism remains operational under various environmental conditions, albeit with varying degrees of efficiency.

## Results

### *CUC2* is derepressed in *clf-81 sep3-2* but is post-transcriptionally dampened

In Arabidopsis, multiple PRC2 catalyze the repressive H3K27me3 chromatin modification (Förderer et al., 2016; Shu et al., 2020; Xiao and Wagner, 2015). There are at least three PRC2 complexes, each containing one of the 3 histone methlytransferase CURLY LEAF (CLF), SWINGER (SWN) or MEDEA (MEA). Whereas MEA function is restricted to gametogenesis and endosperm development, CLF has a prominent function during vegetative development, with SWN playing redundant roles (Chanvivattana et al., 2004). Weak mutant lines affected in different PRC2 components show increased leaf dissection (Lafos et al., 2011; Lopez-Vernaza et al., 2012; Müller-Xing et al., 2014). However, strong PRC2 mutants such as strong *clf* mutants have very pleiotropic effects, forming small rosettes with small narrow, curled leaves. In addition few leaves are produced as plants are very early flowering (Goodrich et al., 1997; Lopez-Vernaza et al., 2012; Schubert et al., 2006), which severely hampers the analysis of leaf shape. This strong phenotype is in part due to derepression of floral genes during the vegetative phase (Goodrich et al., 1997) and part of the *clf* phenotype is therefore the secondary effect of a change in the plant developmental program. To limit these effects we analysed the *clf-81 sep3-2* double mutant that combines the strong *clf-81* allele, containing a missense mutation of a conserved amino acid residue within the catalytic SET domain (Schubert et al., 2006), with a mutation in the class E floral homeotic gene *SEP3* that limits the expression of floral genes in vegetative tissues and partially restores normal flowering time (Lopez-Vernaza et al., 2012). Therefore, the *clf-81 sep3-2* makes it possible to analyze the effects of a strong *CLF* mutation independently of most of its secondary effects (Figure S1). Morphometric analyses using MorphoLeaf (Biot et al., 2016; Oughou et al., 2023) showed that leaves of *clf-81 sep3-2* mutants grown under short-day (SD) were more dissected than wild type with a higher LDI and TAR (Figure 1b, c).

The *CUC2* gene is targeted by PRC2 and marked by H3K27me3 and CLF/SWN occupancy in different tissues (Bouyer et al., 2011; Lafos et al., 2011; Roudier et al., 2011; Shu et al., 2019). We confirmed by ChIP followed by qPCR analyzes that the *CUC2* locus was strongly marked by H3K27me3 and bound by CLF in tissues enriched in young developing leaves (Figure 1 e,f, Figure S2a-c). To further test H3K27me3 deposition in leaf sinus cells expressing CUC2, we isolated such cells using an INTACT approach and found that they also showed H3K27me3 deposition on *CUC2* (Figure 1g). This suggests that PRC2 may regulate *CUC2* transcription. To test this, we introduced the pCUC2:RFP transcriptional reporter in *clf-81 sep3-2* and quantified fluorescence intensity in the distal sinus of tooth 1 in two classes of leaf blade length (250 to 750 µm and 750 to 1250 µm), that correspond respectively to the early and late stages of primary leaf morphogenesis. pCUC2:RFP reporter is expressed at higher levels in the sinus of *clf-81 sep3-2* compared to WT (Figure 1h), indicating that *CUC2* is subject to negative transcriptional regulation by PRC2 in the leaf sinuses. We next introduced the translational pCUC2:CUC2-VENUS reporter into *clf-81 sep3-2* to quantify the dynamics of CUC2 protein accumulation. Surprisingly, CUC2-VENUS fluorescence is not increased in the *clf-81 sep3-2* sinus compared to WT (Figure 1i). Next, to identify the developmental origin of the increased *clf-81 sep3-2* dissection, we retraced its developmental trajectory from static images using MorphoLeaf (Biot et al., 2016; Oughou et al., 2023). This analysis showed that *clf-81 sep3-2* leaves become more dissected than WT late in development during secondary morphogenesis, once the leaf blade is longer than 1000 µm (Figure S3). In contrast, during the primary morphogenesis phase, *clf-81 sep3-2* teeth 1 have similar shapes and the primordia similar LDI compared to WT (Figure 1j,k), in agreement with previous analysis (Oughou et al., 2023). Therefore, both leaf morphometrics and CUC2 protein quantification indicated that increased *clf-81 sep3-2* leaf dissection occurs during secondary morphogenesis, which can not be related to an increased CUC2 protein level. Altogether, these observations indicate that in *clf-81 sep3-*2 a transcriptional derepression of *CUC2* occurs during primary morphogenesis but that a stronger post-transcriptional inhibition dampens CUC2 protein levels and maintains unchanged primary morphogenesis.

### MIR164A activity is not increased in *clf-81 sep3-2*

Because *MIR164A* regulates *CUC2* expression during the leaf development (Maugarny-Calès et al., 2019; Nikovics et al., 2006) we hypothesized that post-transcriptional dampening of CUC2 in *clf-81 sep3-*2 could result from an increased activity of *MIR164A*. This hypothesis led to 3 testable predictions. First, the expression level of *MIR164A* should be increased in the *clf-81 sep3-2* background compared to WT. To test this, we introduced the pMIR164A:RFP reporter in *clf-81 sep3-2*. Quantification of pMIR164A:RFP fluorescence did not reveal any increase in *clf-81 sep3-2* compared to WT, indicating that pMIR164A is not more expressed (Figure 2a). Second, higher CUC2 protein levels should be observed in *clf-81 sep3-2 mir164a-4* compared to *mir164a-4*. However, quantification of pCUC2:CUC2-VENUS fluorescence did not reveal any increase in *clf-81 sep3-2 mir164a-4* compared to *mir164a-4* (Figure 2b). Third, young leaf primordia of *clf-81 sep3-2 mir164a-4* should be more dissected than those of *mir164a-4*. Morphometric analysis did not reveal any increase in leaf primordium dissection in *clf-81 sep3-2 mir164a-4* compared to *mir164a-4*, and our quantifications rather pointed to a reduced tooth pointiness and lower LDI (Figure 2c, d). In conclusion, quantification of the *MIR164A* promoter activity, CUC2 protein levels or leaf morphology converge to indicate that *MIR164A* activity is not increased in *clf-81 sep3-2* compared to WT and that *MIR164A* is not responsible for CUC2 dampening in *clf-81 sep3-2*.

**Figure 2:**
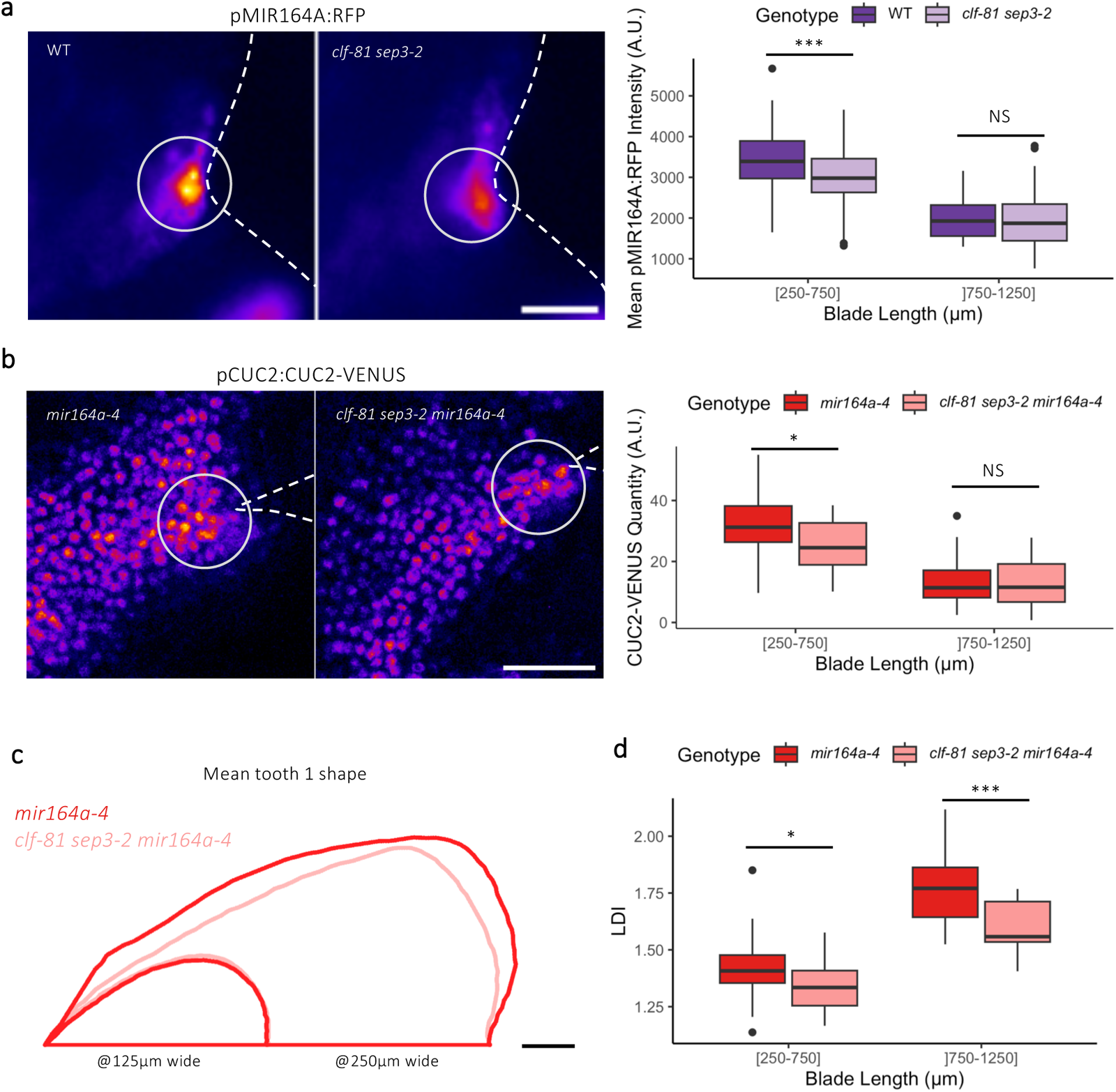
*MIR164A* is not responsible for CUC2 dampening in the *clf81 sep3-2* mutant. a) pMIR164A:RFP intensities in WT and *clf-81 sep3-2* distal sinus of tooth 1. b*) pCUC2:CUC2-VENUS* intensities in *mir164a-4* and *clf-81 sep3-2 mir164a-* distal sinus of tooth 1. a) and b) Images on the left are representative of the typical relative intensities obtained in the different genotypes. Dashed lines mark the leaf margin limit, the grey circles the quantification areas. Quantifications are plotted in arbitrary unit (A.U.) for two bins of blade length, *n* ≥ 39 for a) and *n* ≥ 19 for b). c) Mean shape of tooth 1 in *mir164a-4* and *clf-81 sep3-2 mir164a-4* developing leaves. Small and large teeth are 125µm and 250 µm wide, respectively, which correspond to leaf primordia about 500µm and 1000µm long. d) LDI of *mir164a-4* and *clf-81 sep3-2 mir164a-4* developing leaves *n* ≥ 17. Statistical significance is tested by Student tests: NS for not significant, p-value is <0.05 for *, <0.01 for **, <0.005 for ***. Scale bars in a), b) and c) is 30 µm.

### *MIR164B* is targeted by CLF and contributes to CUC2 dampening in *clf-81 sep3-2*

Since our analysis excluded *MIR164A* as responsible for CUC2 dampening in *clf-81 sep3-2* mutants, we next turned to the two other *MIR164* genes, *MIR164B* and *MIR164C*. Because floral genes are ectopically expressed in l *clf* eaves and *MIR164C* has a well-described role in floral development, *MIR164C* was a good candidate (Baker et al., 2005; Goodrich et al., 1997; Sieber et al., 2007). To test the potential involvement of *MIR164B* or *MIR164C* in CUC2 dampening we first analyzed whether they were regulated by PRC2. The repressive H3K27me3 mark are present on both *MIR164B* and *MIR164C* in both young developing leaf tissues and CUC2-expressing cells (Figure 3 a, b, d, Figure S2d-f). In contrast, CLF ChIP-qPCR showed that CLF significantly binds to the *MIR164B* locus and only weakly to *MIR164C* (Figure 3c).

**Figure 3:**
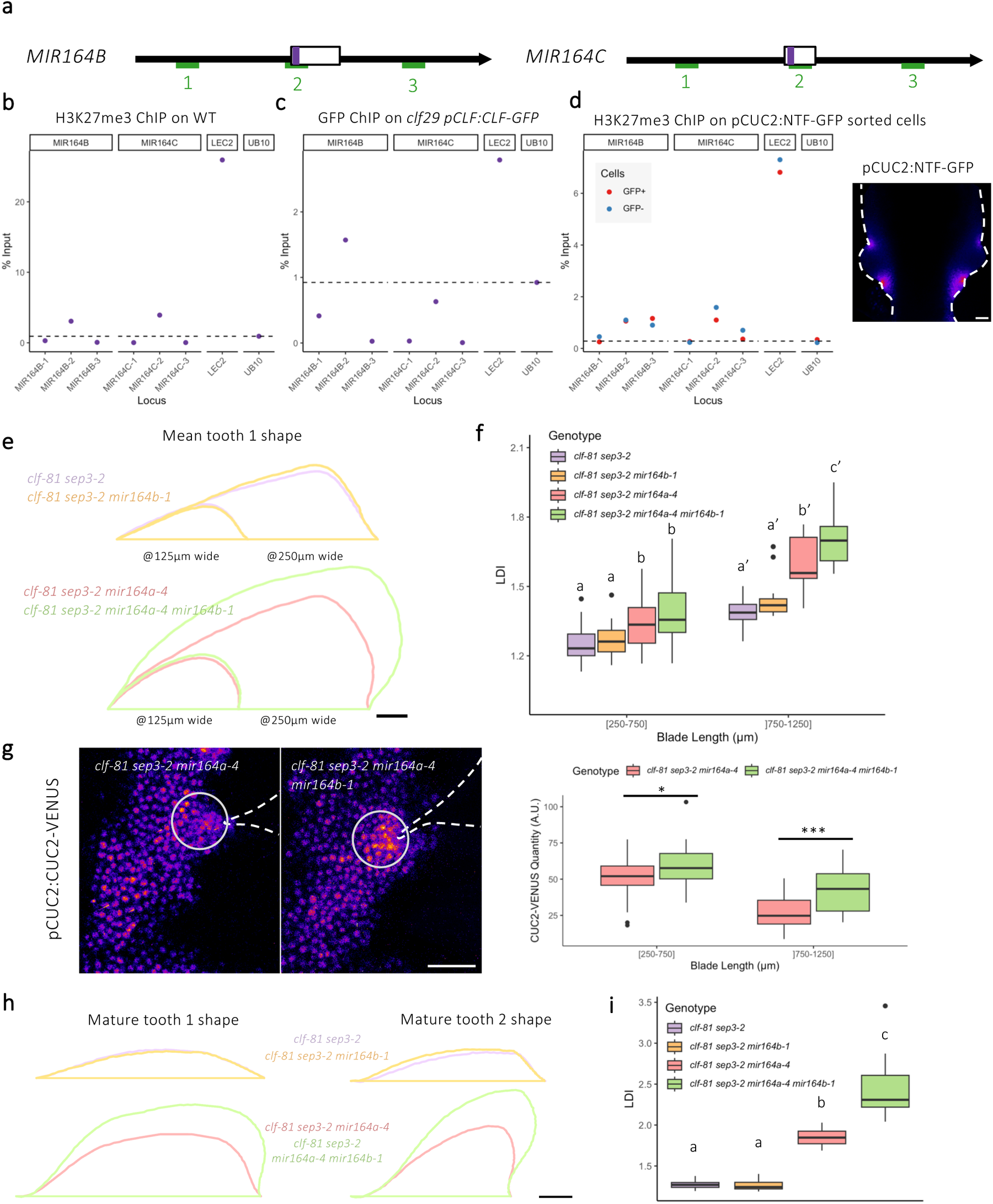
*MIR164B* is targeted by CLF and reduces CUC2 protein levels in the *clf81 sep3-2* mutant. a) Schematic *MIR164B* and *MIR164C* gene structures. The primary miRNA sequence is represented by a box and the mature miRNA sequence is highlighted in purple. The amplified regions for ChIP-qPCRs in b), c) and d) are defined by the three sets of primers represented in green. b) H3K27me3 chromatin-immuno-precipitation followed by qPCR (ChIP-qPCR) on WT rosette centers. c) GFP ChIP-qPCR on *clf29* pCLF:CLF-GFP rosette centers. d) H3K27me3 ChIP-qPCR *on pCUC2:NTF-GFP* rosette centers GFP sorted nuclei. In d), the picture shows the expression pattern in a young leaf of the pCUC2:NTF-GFP reporter which was used to sort out GFP+ and GFP-cells. Dashed lines mark the leaf margin. The results of a single experiment are shown, the results of other biological replicates are available in Figure S2d-e. The *LEC2* locus is known to be marked by H3K27me3 in this type of tissue (positive control), while the *UB10* locus acts as a negative control. (e) Mean shape of tooth 1 in *clf81 sep3-2*, *clf81 sep3-2 mir164b-1*, *clf81 sep3-2 mir164a-4* and *clf81 sep3-2 mir164a-4 mir164b-1* developing leaves. Small and large teeth are 125µm and 250 µm wide, respectively, which correspond to leaf primordia about 500µm and 1000µm long. f) LDI of in *clf81 sep3-2*, *clf81 sep3-2 mir164b-1*, *clf81 sep3-2 mir164a-4* and *clf81 sep3-2 mir164a-4 mir164b-1* developing leaves. *n* ≥ 15. g) pCUC2:CUC2-VENUS intensities in *clf81 sep3-2 mir164a-4* and *clf81 sep3-2 mir164a-4 mir164b-1* distal sinus of tooth 1. Images on the left are representative of the typical relative intensities. Dashed lines mark the leaf margin limit, the grey circles the quantification areas. Quantifications are plotted in arbitrary unit (A.U.) for two bins of blade length, *n* ≥ 23. h) Mean shape of tooth 1 and tooth 2 from *clf81 sep3-2*, *clf81 sep3-2 mir164b-1*, *clf81 sep3-2 mir164a-4* and *clf81 sep3-2 mir164a-4 mir164b-1* mature leaves. i) Mature LDI of *clf81 sep3-2*, *clf81 sep3-2 mir164b-1*, *clf81 sep3-2 mir164a-4* and *clf81 sep3-2 mir164a-4 mir164b-1*. *n* ≥ 22. Statistical significance is tested by Student’s tests in g) (NS for ‘not significant’, p-value is <0.05 for *, <0.01 for **, <0.005 for ***) and ANOVA followed by Tukey HSD in f) and i). ANOVA are performed within leaf blade bins in f). Scale bars in d), e) and g) are 30 µm and in h) 1 mm.

This prompted us to test whether *MIR164B* regulated CUC2 when PRC2 activity was compromised. For this, we analyzed early leaf morphogenesis in *clf-81 sep3-*2 mutants in which *MIR164B* was mutated (together with *MIR164A* or not) (Figure 3 e, f). Thus, while tooth shape and LDI are identical for *clf-81 sep3-2 mir164b-1* and *clf-81 sep3-2,* we observed that teeth are more pronounced and the LDI higher in the 750-1250 µm class of *clf-81 sep3-2 mir164a-4 mir164b-1* compared to *clf-81 sep3-2 mir164a-4* (Figure 3 e, f). We did not observe any significant difference for these two genotypes in the smaller 250-750 µm class. Together, these morphometric analyses pointed to a role of *MIR164B* during late primary morphogenesis in *clf-81 sep3-2,* a role which was masked by *MIR164A.* To confirm this, we compared the fluorescence of the pCUC2:CUC2-VENUS reporter in *clf-81 sep3-2 mir164a-4 mir164b-1* and *clf-81 sep3-2 mir164a-4.* This pointed to higher CUC2 levels in *clf-81 sep3-2 mir164a-4 mir164b-1* compared to *clf-81 sep3-2 mir164a-4,* which was more pronounced in the *750-1250µm* class compared to the *250-750µm* class (Figure 3g). Therefore, we concluded that *MIR164B* contributes to dampen CUC2 protein levels to control late primary morphogenesis when CUC2 promoter activity is increased in *clf-81 sep3-2*. Importantly, such an effect of MIR164B on primary morphogenesis is not observed in a wild-type *CLF* background, either in presence or absence of *MIR164A* (Figure S3 a, b), indicating a direct or indirect relationship between the action of MIR164B and PRC2 activity at the CUC2 locus.

The effect of *MIR164B* on leaf shape is also visible at the mature leaf stage, as *clf-81 sep3-2 mir164a-4 mir164b-1* have more pronounced teeth and higher LDI compared to *clf-81 sep3-2 mir164a-4* (Figure 3 h, i). Again, such contribution of *MIR164B* on mature leaf shape is not observed in genotypes with functional CLF (Figure S3 c, d). Finally, the contribution of *MIR164B* on leaf morphology is not dependent on the *sep3-2* mutation nor specific to the *clf-81* allele as it is also observed for the *clf-29* allele, though with a milder effect, possibly because *clf-29* leads to a weaker phenotype compared to *clf-81* (Figure S3 e, f).

Altogether, this led us to propose the following model (Figure 4a). Compromised PRC2 activity leads to increased secondary morphogenesis independently of CUC2. In parallel, it leads to CUC2 and *MIR164B* transcriptional derepression, which compensate each other to provide wild-type CUC2 protein levels and primary morphogenesis. When *MIR164B* is simultaneously impaired by mutation, compromised CLF functions leads to higher CUC2 protein levels and more dissected leaves at the end of the primary morphogenesis. Because such a contribution of *MIR164B* on early morphogenesis was not observed in a CLF+ background, we suggest that such a role of MIR164B results from its derepression in a background compromised for PRC2. This generates a MIR164B-dependant rescue mechanism that provides robustness to CUC2 protein levels and primary leaf morphogenesis following compromised PRC2.

**Figure 4:**
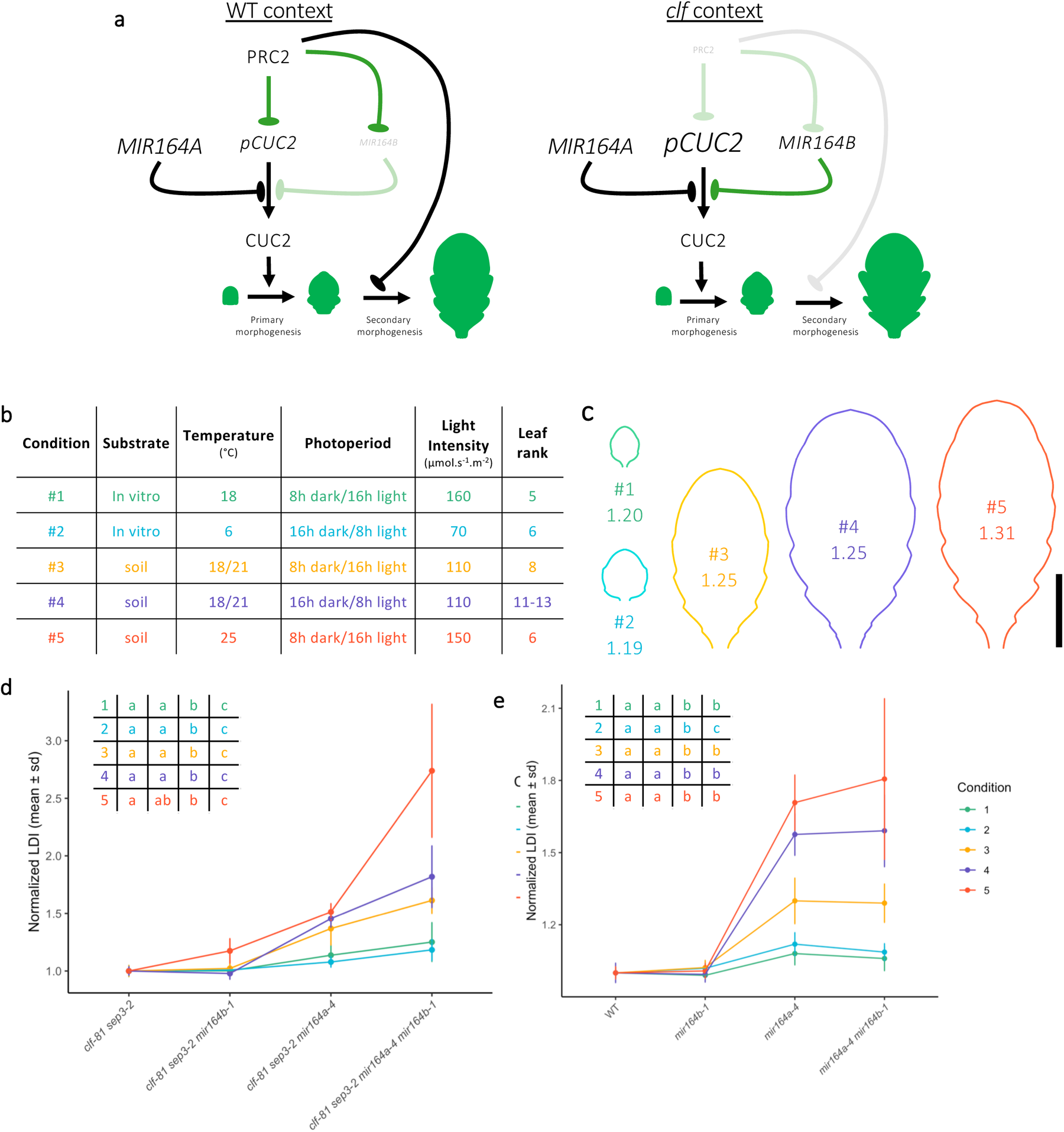
*MIR164B* compensation of defective PRC2 function provides robustness to leaf development under different environmental conditions. a) Genetic model summarizing the interactions between *CUC2* promoter activity (*pCUC2*), CUC2 protein levels (CUC2), *MIR164A*, *MIR164B*, PRC2 and how these actors affect mature leaf dissection. Pointy arrows designate promoting interactions while oval arrows represent inhibitory interactions. Green lines form a feed forward loop, which provides genetic robustness to the CUC2 protein level. In the wild type, CUC2 protein level is transcriptionally regulated by PRC2 (CLF) and post-transcriptionally by *MIR164A*. CUC2 protein levels determine tooth formation during primary morphogenesis. Mature leaf shape is acquired during secondary morphogenesis which is regulated by PRC2 (CLF), independently of CUC2. In the PRC2 (*clf*) mutant background, *CUC2* is transcriptionally derepressed, but CUC2 protein levels are unchanged through the additional action of *MIR164B* which is simultaneously derepressed. As a result, primary morphogenesis is not modified but secondary morphogenesis is modified by the effects on unknow PRC2 (CLF) targets, resulting in more dissected mature leaves. b) Table summarizing the different environmental conditions used to challenge the MI164B-dependant compensation mechanism. Conditions #4 is the one which was used in the experiments described in Figure 1 to 3. c) Average shapes of WT leaves grown in the conditions described in b). d) and e) Mean normalized mature LDI in *clf-81 sep3-2*, *clf81 sep3-2 mir164b-1*, *clf-81 sep3-2 mir164a-4* and *clf-81 sep3-2 mir164a-4 mir164b-1* mutants in d) and WT, *mir164b-1*, *mir164a-4* and *mir164a-4 mir164b-1* in e). LDI was normalized respectively by *clf-81 sep3-2* mean LDI in d) and WT mean LDI in e). Statistical significance is tested by ANOVA followed by Tukey HSD in d) and e) within each environmental condition. Statistical significance is reported in the table on the upper left corner of the plot, in the corresponding order of the genotypes. Error bars are SD, *n* ≥ 4 for d) and 5 for e). Scale bar in c) is 1 cm.

### *MIR164B* compensation of defective PRC2 function is active under different environmental conditions

Genetic regulatory networks controlling leaf shape are modified in response to environmental conditions to provide leaf shape plasticity (Chitwood and Sinha, 2016). Therefore, we wanted to test whether the *MIR164B* compensation mechanism was activated in response to *PRC2* impaired function under different environmental conditions. Thus, we analyzed leaf dissection in the same eight mutant combinations as used before grown under 4 novel environmental conditions (in addition to condition #4 used in the previous experiments). These included variations in growth substrate, temperature, light regime and intensity, leaf rank (Figure 4b). Under these conditions, WT leaf morphology showed very high variations, with modification in its size, overall shape and level of dissection (Figure 4c). Next, we measured the LDI for each genotype under the different conditions. To allow comparison under different environmental conditions, we normalized the LDI and TAR of teeth 1 and 2 to the WT value for all the genotypes in which CLF and SEP3 were active and to the value of *clf-81 sep3-2* for the genotypes mutated for *clf* and *sep3* (Figure 4d,e, Figure S5) and computed average tooth 1 and 2 shapes (Figure S5). This showed that under all tested conditions the normalised LDI and TARs are significantly higher in *clf-81 sep3-2 mir164a-4 mir164b-1* compared to *clf-81 sep3-2 mir164a-4* and similar between *clf-81 sep3-2 mir164b-1* and *clf-81 sep3-2* (Figure 4d, Figure S5a,c, Figure S6). In contrast, in the backgrounds where CLF was active, the mutation of *MIR164B* did not increase leaf dissection, whether *MIR164A* was active or mutated (Figure 4e, Figure S5b, d, Figure 6). This indicated that the *MIR164B*-dependent compensation mechanism was active in all environmental conditions following compromised PRC2 function. However, the increase in dissection observed in *clf-81 sep3-2 mir164a-4 mir164b-1* depended on the growth conditions, being very high in conditions #5, suggesting a strong requirement for the compensation mechanism under elevated temperature. The increased dissection in *clf-81 sep3-2 mir164a-4 mir164b-1* appeared also more pronounced for plants grown in soil compared to plants grown *in vitro*. From this combinatorial analysis, we concluded that the *MIR164B-dependent* compensation mechanism is active under all environmental conditions tested, though with a varying strength.

## Discussion

Here, combining morphometrics, *in situ* gene expression quantification and epigenetic characterisations, we show that *MIR164B* provides robustness to CUC2 protein levels and primary leaf morphogenesis despite higher CUC2 promoter activity due to PRC2 deficiency. We propose that reduction of PRC2 activity simultaneously derepresses *CUC2* and *MIR164B* expression (Figure 4a). In this view, derepressed MIR164B would either lead to a higher expression level or a shift in its expression pattern allowing it to now significantly regulate *CUC2*. Thus, inactivation of PRC2 activates a novel regulatory edge involving *MIR164B* which acts additively to *MIR164A* and provides genetic canalization to variations in *CUC2* expression. This example of epigenetic derepression of miRNA genes enabling developmental robustness is similar to a mechanism of transposon silencing involving miRNAs and secondary siRNAs called Epigenetically activated small interfering RNAs (easiRNAs) (Creasey et al., 2014). In this case, reduced DNA methylation leads simultaneously to the transcriptional derepression of transposons and miRNAs that indirectly lead to the production of easiRNAs which in turn silence the reactivated transposons. Further studies will be required to determine in which context decreased PRC2 activity occurs during development and activates the MIR164B rescue mechanism. Finally, we provide evidence that the increased dissection observed in PRC2 mutant mature leaves occurs during the secondary morphogenesis phase independently of CUC2 through the mis-regulation of other PRC2 targets that need to be identified.

Our study shows that MIR164B provides genetic robustness to leaf morphogenesis, which echoes studies in the animal development pointing to miRNAs as major actors of genetic robustness (Alberti and Cochella, 2017; Avital et al., 2018; Hornstein and Shomron, 2006; Posadas and Carthew, 2014). Two scenarios have been described in this case. First, a regulatory gene is activating the expression of a miRNA and repressing at the same time the miRNA target, forming a coherent feed forward loop (FFL) in which the regulatory gene and the miRNA combine their effects to repress the target. Such circuits have been associated with clearing of target gene expression to maintain sharply contrasted expression domains. Our model does not fit with this case, but with the second case, in which the regulatory gene has the same effect (activating or repressing) on both the miRNA and its target, thus forming an incoherent FFL. Here, PRC2 represses both *CUC2* and *MIR164B* forming an incoherent FFL of type 2 (Alon, 2007). Such incoherent networks have been shown to allow robust target gene expression in face of environmental variation or genetic noise resulting from stochasticity in transcription (Li et al., 2009). In our system the target CUC2 does not experience noisy expression but increased expression due to epigenetic derepression. The predicted outcome of such networks is to stabilize target gene level, here CUC2, which is what we observe when we quantify CUC2 protein levels and its morphological effect. Like in animals in which the phenotype of mutants in the miRNA contributing to robustness may only appear under external stresses (Cassidy et al., 2013; Li et al., 2009), here the role of miR164B is only revealed under a genetic stress induced by compromised PRC2 function.

*MiR164* is encoded by 3 genes, two of them MIR164A and MIR164C being expressed respectively in leaf and floral domains overlapping with those of their targets CUC1 and/or CUC2 where they control their expression level, without significantly affecting their expression domain (Baker et al., 2005; Nikovics et al., 2006). However, randomly located *CUC2* expression domains were occasionally observed in a triple *mir164abc* mutant, suggesting that collectively the 3 *MIR164* genes may also contribute to robustness of their target expression patterns (Sieber et al., 2007). Nevertheless, the role of individual *MIR164* genes in robustness was not established and, more precisely, the role of *MIR164B* during plant development was not known, though MIR164B was shown to be involved in the physiological response of plants to environmental stresses, but possibly through targets different from CUC1/CUC2 (Du et al., 2022; Tsai et al., 2023). Here we show that *MIR164B* acts as a safe guard to counteract transcriptional deregulation of its target CUC2.

Within the many factors affecting CUC2 expression that have been reported (Maugarny et al., 2015) some may have a similar role as *MIR164B* in providing robustness to CUC2 expression. NGATHA-like transcription factors act as negative regulators of *CUC* expression and are expressed in mostly overlapping expression domains in various plant organs (Engelhorn et al., 2012; Nicolas et al., 2022; Shao et al., 2020; Zhang et al., 2015). Interestingly, their mutation leads to stochasticity in the development of axillary meristems which is suppressed if CUC2 is inactivated (Nicolas et al., 2022), suggesting that CUC2 is causing such lack of developmental robustness. CUC2 expression is also tightly connected with auxin signalling with which it forms a negative feedback loop required for the robust formation of alternate patterns of high CUC2 or high auxin patterns at the leaf margin (Bilsborough et al., 2011). All these observations point to a highly connected CUC2 regulatory module which could form the core network of a so-called bow-tie structure. Such bow-tie structures are formed by a central, dense, highly connected network in which many different inputs are feeding in and which feeds to many downstream components. Such regulatory modules have been suggested to contribute to the robustness of many different biological systems (Kitano, 2004; Whitacre, 2012). In such a view, modulation of the “CUC2-knot” of the bow-tie by environmental factors could contribute to leaf shape plasticity. Our observations indicate that the MIR164B node itself may be modulated by environmental conditions. Therefore, further studies will be required to determine if leaf plasticity involves modulation of the CUC2 regulatory network and if this occurs through modulation of factors contributing to its robustness such as MIR164B.

## Material and Methods

### Plant material

The lines *clf-81 sep3-2* (Lopez-Vernaza et al., 2012), *clf-29* (Bouveret et al., 2006), *mir164a-4* (Nikovics et al., 2006), *mir164b-1* (Mallory et al., 2004), pCUC2:RFP (Gonçalves et al., 2015), *pMIR164:RFP* (Maugarny-Calès et al., 2019), the complemented *pCUC2:CUC2-VENUS cuc2-3* (Gonçalves et al., 2015) and the complemented *pCLF:CLF-GFP* genomic fusion in *clf-29* (de Lucas et al., 2016) are in a Col-0 background. All multiple mutants were generated by crossings, verified by genotyping and maintained as multiple homozygous lines. Note that when the *pCUC2:CUC2-VENUS* reporter was used it was associated with the *cuc2-3* mutant background: for instance, the *clf-81 sep3-2 pCUC2:CUC2-VENUS* line used in Figure 1i was also homozygous mutant for *cuc2-3*.

### Cloning and transgenic plants selection

The promoter of the *CUC2* gene was amplified with primers GGGGACAACTTTGTATAGAAAAGTTGGCACCTCCTTCATCAAATACG and GGGGACTGCTTTTTTGTACAAACTTGGAAAGATCTAAAGCTTTTGTTTGAGAG and cloned into the entry vector pENTR5’ (Life Technologies). The promoter of CUC2 was fused to a Nuclear Tagging Fusion (NTF) containing the GFP and a nuclear envelop-targetting domain, described in (Morao et al., 2018)) into the pB7m24GW, 3 vector (VIB-UGENT, Karimi et al., 2002) using Gateway technology and following the protocol of the LR Clonase II Plus Enzyme Mix (ThermoFischer). The final pCUC2:NTF-GFP vector was used to transform plant by floral dipping (Clough and Bent, 1998). Transformed plants were selected on basta (10 mg/L) in MS medium from Duchefa. Plants expressing GFP at the expected CUC2 domain were selected after confocal microscopy imaging (LSM 700 Laser-scanning confocal microscope, Zeiss).

### Growth conditions

Growth conditions are indicated in Figure 4b. For soil grown plants a mix peat/sand (TREF 1018201170, TREF, France), was used as substratum, while Arabidopsis medium from Duchefa was used for *in vitro* grown-plants.

### Leaf imaging for morphometrics

For morphological analysis of young leaf primordia under SD grown conditions, plants were grown for 3 to 4 weeks prior to observations. Leaves of rank 11 to 13 were isolated from the meristem using surgical syringe needles and mounted between slide and coverslip, with the adaxial facing the coverslip. Imaging was done with an Axio Zoom.V16 macroscope (Carl Zeiss Microscopy, Jena, Germany, http://www.zeiss.com/) either in the chlorophyll fluorescence channel using a Zeiss 63 HE filter set (excitation band pass filter 572/25; beam spliter 515, emission band pass filter 535/30) or by transmitted white light. The data set of WT *and clf-81 sep3-2* used for Figures 1j, k and Figure S3 is the same as the one described in Oughou et al., (2023). Mature leaves grown in vitro in conditions #1 and #2 were imaged in the same way. For the observation of mature leaves of soil grown plants, leaves of defined rank (see Figure 4a) were cut with scissors and glued on paper (adaxial side towards the paper) before being scanned in black and white at 1600 dpi with an office scanner (Epson Perfection V800 Photo).

### Reporter imaging and quantification

pCUC2:CUC2-VENUS lines were imaged on a Leica SP5 inverted microscope (Leica Microsystems, Wetzlar, Germany). Lenses are Leica 20x HCX PL APO CS. pCUC2:CUC2-VENUS was excited at 514 nm and fluorescence was collected between 530 and 580 nm. Acquisition parameters were kept constant throughout acquisitions so that intensity levels are comparable. Signal was quantified on the sinus distal to the first tooth, as *CUC* expression in this domain has been shown to drive the outgrowth of marginal structures (Abley et al., 2016; Blein et al., 2008). The intensity of the 12 most intense nuclei was measured manually using Fiji on the medial plane of each nucleus. The mean intensity of the background was substracted from the mean of the intensity of the nuclei.

pCUC2:RFP and pMIR164A:RFP lines were imaged on an Axio Zoom.V16 macroscope (Carl Zeiss Microscopy, Jena, Germany, http://www.zeiss.com/), using a custom-made filter block (excitation band pass filter 560/25; beam spliter 585, emission band pass filter 615/24, AHF, Tuebingen, Germany, https://www.ahf.de/). Using Fiji, the distal sinus region was cropped from the leaf pictures and the mean intensity in the brightest 500µm^2^ region (which correspond to about 12 cells) was measured using the Qpixie Fiji macro previously described (Gonçalves et al., 2017). All reporter images are displayed using the LUT ‘Fire’ from Fiji.

### Morphometric analyses and mean tooth shape modelling

All leaf shape quantifications and modelling of mean shapes were done using MorphoLeaf software that runs of free-D (Andrey and Maurin, 2005; Biot et al., 2016; Oughou et al., 2023). Briefly, stacks of leaf pictures of the same condition are generated, the leaf contour is automatically segmented and hand-corrected if required. The junctions between the blade and petiole are marked manually and the tip of leaf, tooth sinuses, tooth peaks are automatically extracted, with a manual correction step. From the leaf contour and landmarks, blade length, tooth width, tooth height, blade perimeter and area, alpha shape perimeter and area are computed, and TAR and LDI are calculated as indicated in Figure 1a.

Mean tooth shape modelling of developing leaves was done as follows. First a series of mean leaf shapes of increasing blade length was generated using the “sliding average” function of MorphoLeaf at a leaf blade step size of about 3-5µm. Then, the width of tooth 1 was calculated for each of the resulting mean leaf shapes and the tooth which width was closest to 125 or 250µm was extracted to provide the model of the mean tooth at the expected size. Teeth of 125µm or 250µm width were selected because they roughly lie on leaf primordia with a blade of 500µm and 1000µm long (Table S1), which correspond to the median sizes of the early and late primary morphogenesis phases we defined (respectively intervals of 250 to 750µm and 750 to 1250µm).

Mean tooth shape modelling of mature leaves was done as follows. First, the mean leaf shape of each genotype was calculated using the “average based on bins” function of MorphoLeaf. Next the widths of average teeth 1 and 2 were calculated from the average leaf shape. Because of leaf size variability, average leaf blade lengths and tooth widths were not identical for all genotypes. Therefore, to help comparing shapes of teeth independently of small variations in their sizes, we modelled teeth of similar widths. For this we first calculated the mean tooth width for couples of genotypes to be compared (eg WT and *mir164b-1*, Table S2). Next, we computed a series of leaf shapes of increasing blade length using the “sliding average” function of MorphoLeaf at a leaf blade step size of about 50-70 µm. Then, the width of teeth 1 and 2 were calculated for each of the resulting mean leaf shape and the teeth 1 and 2 which width was closest to the mean of the two genotypes previously calculated was extracted to provide the model of the mean tooth at the expected size.

### Morphometric analyses and statistical analysis

All data were analysed using R and details of the statistical test performed are available in Table S3.

### Immuno-affinity purification (INTACT) of CUC2-GFP positive nuclei

INTACT experiments were performed according to (Morao et al., 2018). Briefly, 500 mg of hand-dissected rosette centers from 6-week-old plants were harvested and frozen in liquid nitrogen. After grounding to a fine powder, cross-linking with 1% formaldehyde was performed at room for 7 minutes. Nuclei were isolated using a Dounce tissue grinder. GFP-labelled nuclei were then purified using magnetic beads that were loaded with anti-GFP antibodies (Abcam, Ab290). Multiple washes were performed to remove cellular debris and unlabelled nuclei to obtain a purified GFP+ nuclei suspension. The remaining, GFP negative suspension of nuclei was also recovered for further analyses. Following extraction, chromatin was sonicated using a Covaris S220 ultra-sonicator to generate DNA fragments with an average length of 250 bp. A small proportion of solubilised chromatin was kept as INPUT for each sample, while the rest was incubated overnight with 1 ug of anti-H3K27me3 (Millipore 07-449). Purified DNA fragments were then used for ChIP-qPCR analyses.

### ChIP and qPCR

Chromatin extraction and immunoprecipitation was performed essentially as described in (Bouyer et al., 2018) with minor modifications. Briefly,300 mg of liquid nitrogen-ground, hand-dissected, rosette centers from 5 week-old *clf81 sep3* or 6 week-old WT plants were subject to double crosslinking (2.5 mM di(N-succinimidyl) glutarate 60 min followed by 1% formaldehyde 7 min). Following nuclei purification, chromatin fragmentation was done using a Covaris S220 ultra-sonicator with the following parameters: 12 min with 5% duty cycle, 105W peak power and 200 cycles per burst. Chromatin immunoprecipitation was performed using 3 µg of anti-GFP antibodies (ab290, Abcam) or anti-H3K27me3 (07-449, Merck) pre-bound to protein A/G-coupled magnetic beads (Dynabeads, Invitrogen) (Bouyer et al., 2018). Relative enrichment levels were determined by ChIP-qPCR in an optical 384-well plate in the QuantStudio™ 6 Flex Real-Time PCR System (ThermoFisher Scientific), using FastStart Universal SYBR Green Master (Rox) (Roche), according to the manufacturer’s instructions. The data were analyzed using the QuantStudio Real-Time PCR Software v1.3 (Applied Biosystems). Cp values were converted using the 2-ΔCT method and relative enrichment was expressed as percentage of input. qPCRs were performed in 2 technical replicates for each sample and at least two biological replicates were analyzed independently. Primers used for this assay are given Table S3.

## Authors contribution

Conceptualization, A.M. and P.L.;

Formal Analysis, A.M., A.V. and P.L.;

Investigation, A.M., A.V., B.A., N.M.F. F.R. and P.L.;

Writing – Original Draft, A.M. and P.L.;

Writing – Review & Editing, A.M., A.V., F.R. and P.L.;

Visualization, A.M. and P.L.;

Supervision, F.R. and P.L.;

Funding Acquisition, A.M., F.R. and P.L.

## Acknowledgments

The IJPB benefits from the support of Saclay Plant Sciences-SPS (ANR-17-EUR-0007). We thank members of the FTA team at IJPB for helping sampling tissues for the ChIP experiments. We thank Hervé Frémineur for technical help in leaf contour correction, Eric Biot for help with MorphoLeaf and Dr Pradeep Das for providing the promoter of pCUC2 cloned in the pENTR5’. We thank Dr Hervé Vaucheret for comments on the manuscript and Prof. Justin Goodrich for sharing seeds and comments on the manuscript.

