## Supplementary Figures and Tables for "Compensation of compromised PRC2 regulation by a miRNA ensures robustness of Arabidopsis leaf development"

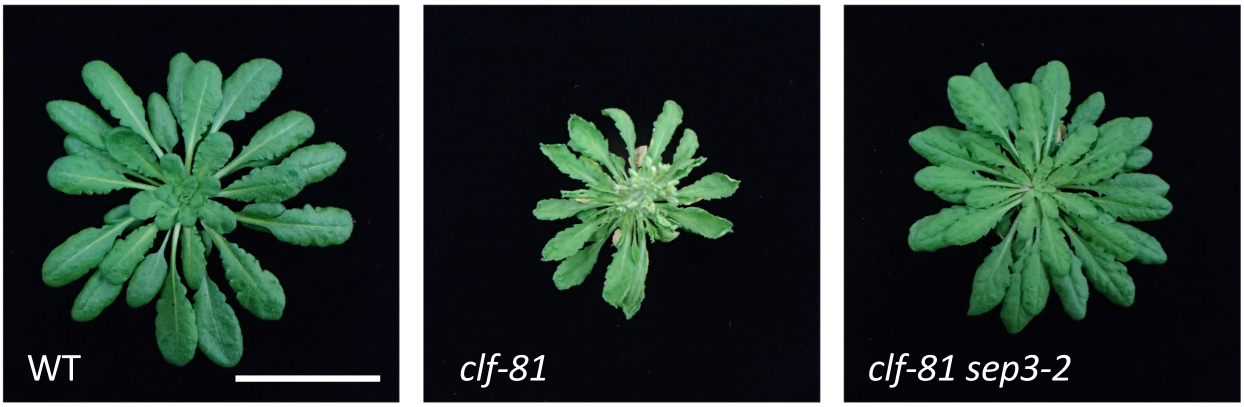


**Figure S1: *sep3-2* partially restores the *clf-81* phenotype.**
Seven week-old rosettes grown in short-day conditions. *clf-81* mutants have very small curled leaves. The *clf-81 sep3-2* double mutant has larger and flat leaves, allowing analyzing the consequence of strong *CLF* mutation independently of the effect of ectopic floral genes expression in leaves.
Scale bar is 5 cm.

Fig S1

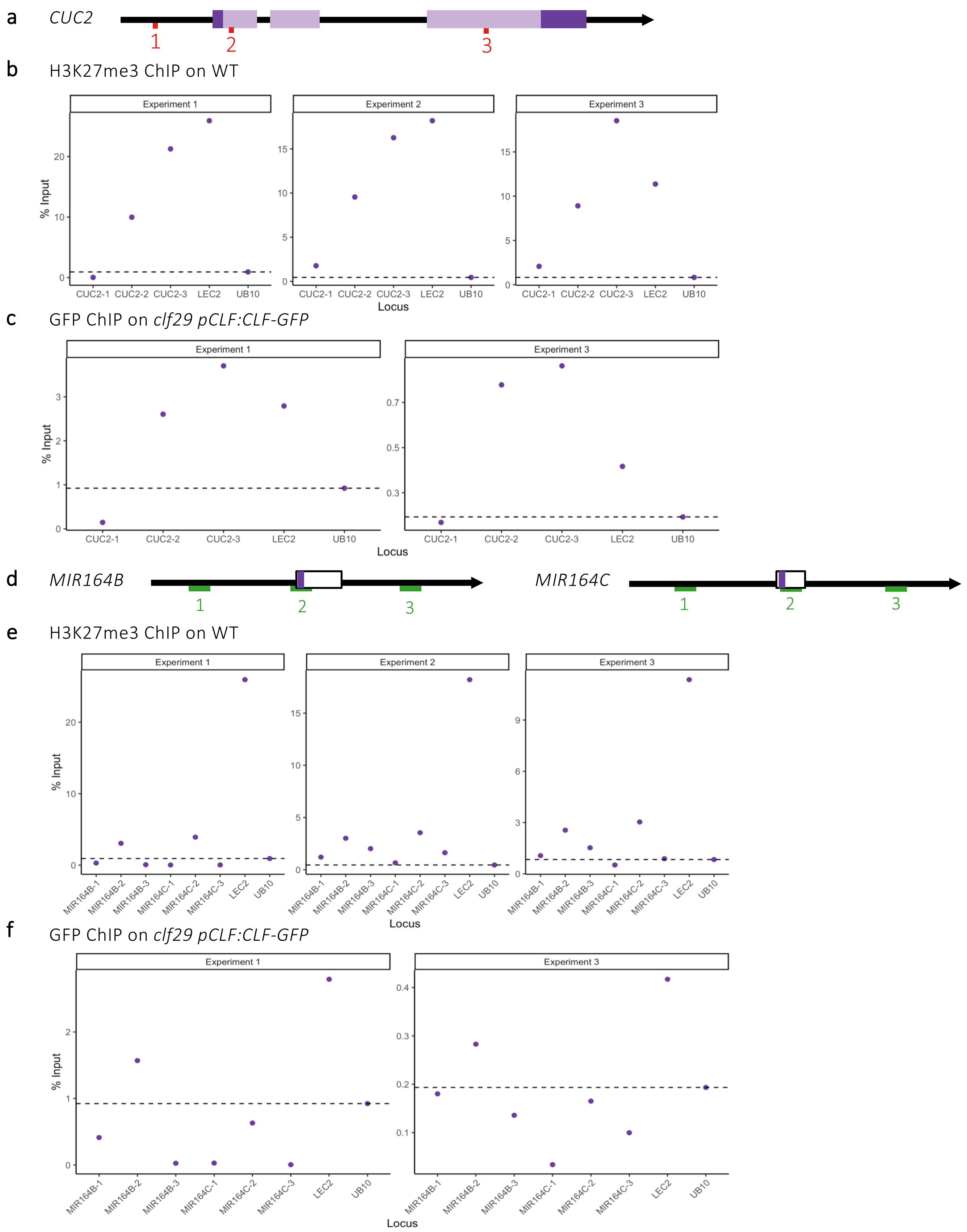


**Figure S2: *H3K27me3 and CLF-GFP ChIP***

a) Schematic *CUC2* gene structure. Exons are represented by light purple boxes (translated regions) and dark purple boxes (untranslated regions), introns and promoter regions are represented by black lines. The amplified regions for ChIP-qPCRs in b) and c) are defined by the three sets of primers represented in green b) H3K27me3 chromatin-immuno-precipitation followed by qPCR (ChIP-qPCR) on WT rosette centers. c) GFP ChIP-qPCR on *clf-29 pCLF:CLF-GFP* rosette centers. d) Schematic *MIR164B* and *MIR164C* gene structures. The primary miRNA sequence is represented by a box and the mature miRNA sequence is highlighted in purple. The amplified regions for ChIP-qPCRs in e) and f) are defined by the three sets of primers represented in green. e) H3K27me3 chromatin-immuno-precipitation followed by qPCR (ChIP-qPCR) on WT rosette centers. f) GFP ChIP-qPCR on *clf29 pCLF:CLF-GFP* rosette centers. The results of a 2 or 3 independent experiment are shown. Results of experiment 1 are shown in Figure 1. The *LEC2* locus is known to be marked by H3K27me3 in this tissue (positive control), while the *UB10* locus acts as a negative control.

Fig S2

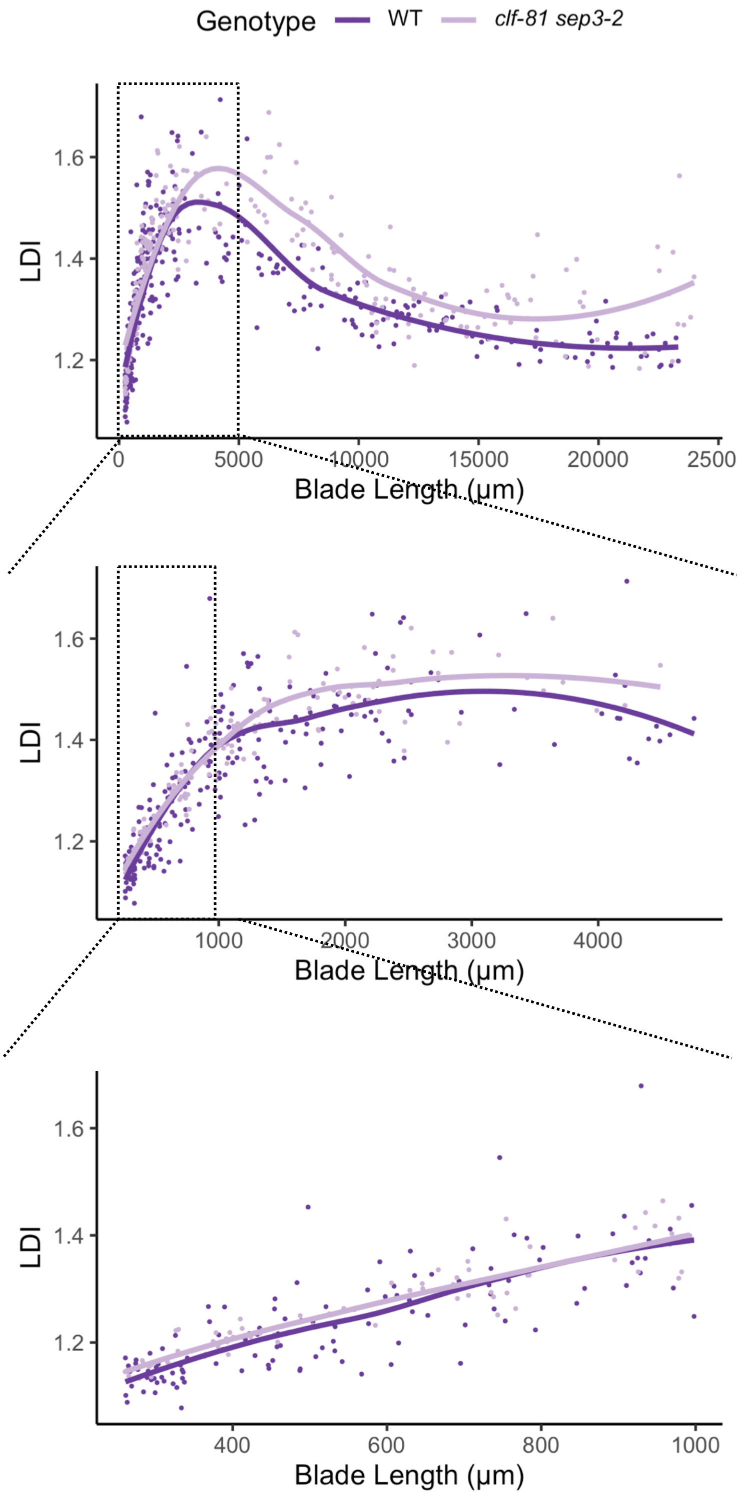


**Figure S3: The increased *clf-81 sep3-2* leaf serration phenotype arises during secondary morphogenesis**

LDI evolution during leaf development. While LDI is higher for *clf-81 sep3-2* compared to WT for leaves larger than 1000µm (secondary morphogenesis), it is similar for smaller leaves that are in the primary morphogenesis phase.

Fig S3

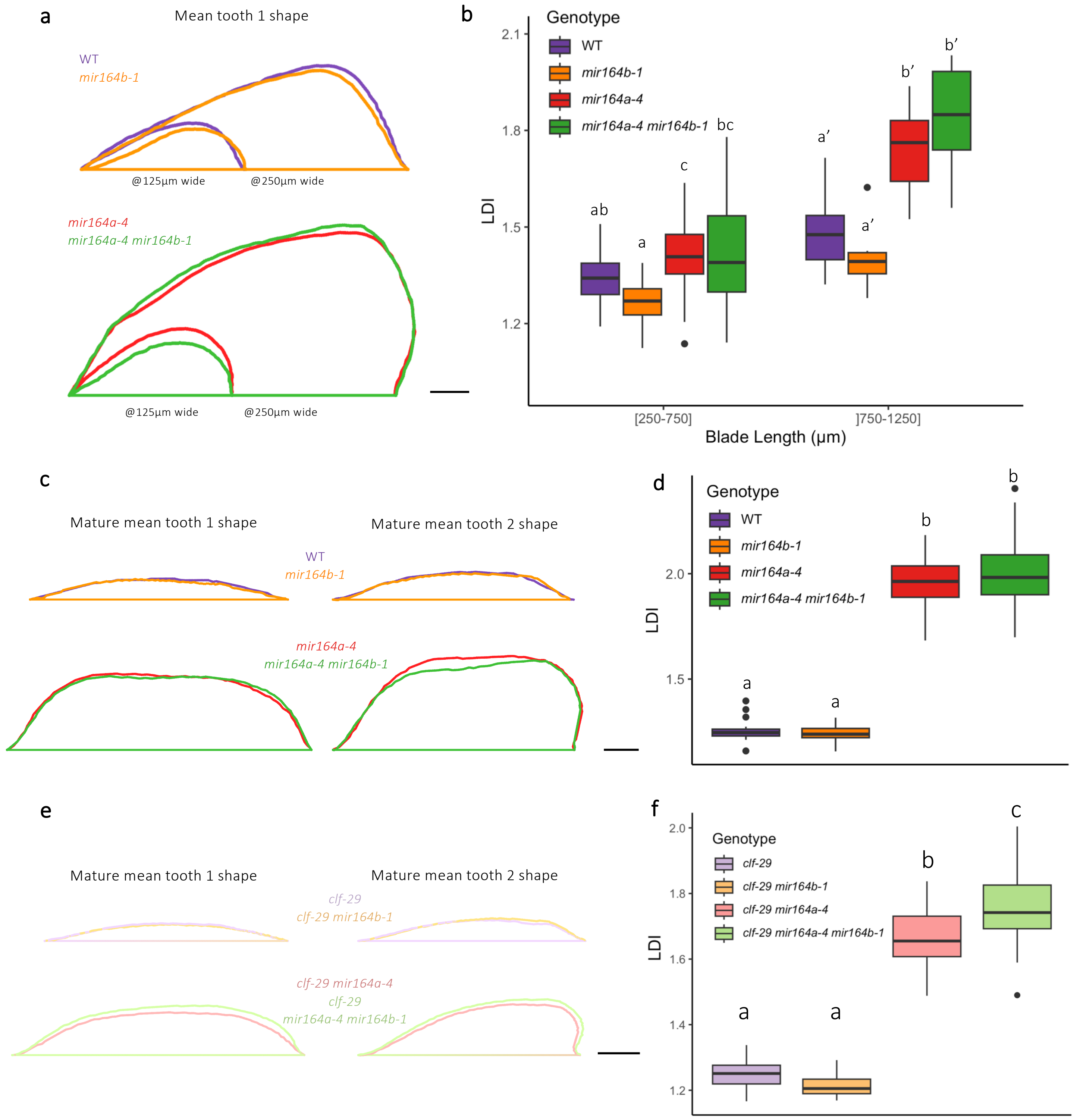


**Figure S4: *MIR164B* mediated rescue mechanism for CUC2 protein levels is active in a *clf-29* mutant background and not in a WT background.**

a) Mean shape of tooth 1 in WT, *mir164b-1*, *mir164a-4* and *mir164a-4 mir164b-1* developing leaves. Small and large teeth are 125µm and 250 µm wide, respectively, which correspond to leaf primordia about 500µm and 1000µm long. b) LDI of WT, *mir164b-1*, *mir164a-4* and *mir164a-4 mir164b-1* developing leaves. $n \geq 12$. c) Mean shape of tooth 1 and tooth 2 from WT, *mir164b-1*, *mir164a-4* and *mir164a-4 mir164b-1* mature leaves. d) Mature LDI of WT, *mir164b-1*, *mir164a-4* and *mir164a-4 mir164b-1*. $n \geq 21$. e) Mean shape of tooth 1 and tooth 2 from *clf-29*, *clf-29 mir164b-1*, *clf-29 mir164a-4* and *clf-29 mir164a-4 mir164b-1* mature leaves. f) Mature LDI of *clf29*, *clf29 mir164b-1*, *clf29 mir164a-4* and *clf29 mir164a-4 mir164b-1*. $n \geq 26$. Statistical significance is tested by ANOVA followed by Tukey HSD in b) d) and e). ANOVA are performed within leaf blade bins in b). Scale bars in a) is 30 µm and in c) and e) 1 mm.

Fig S4

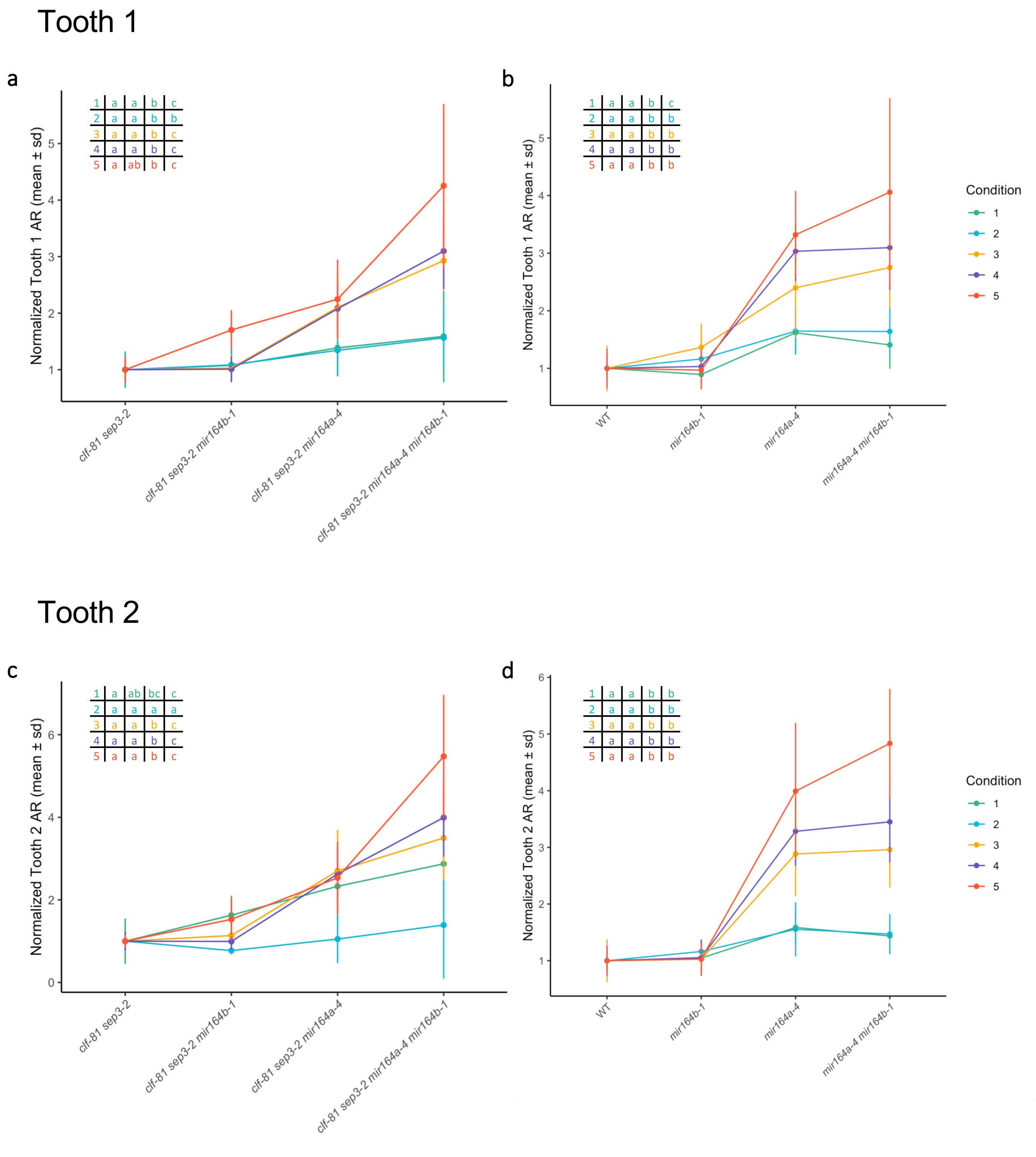


**Figure S5: TAR of tooth 1 and 2 of mature leaves of plants under different growth conditions**

Mean normalized TAR of tooth 1 and 2 of mature leaves of the indicated genotypes under the five different growth conditions (see Figure 4b for details of conditions). TAR was normalized either to *clf-81 sep3-2* (for a and c) or WT (for b and d) mean TAR. Statistical significance is tested by ANOVA followed by Tukey HSD within each environmental condition. Statistical significance is reported in the table on the upper left corner of the plots, in the corresponding order of the genotypes. Error bars are SD, $n \geq 4$ for a) and c) and 5 for b) and d).

Fig S5

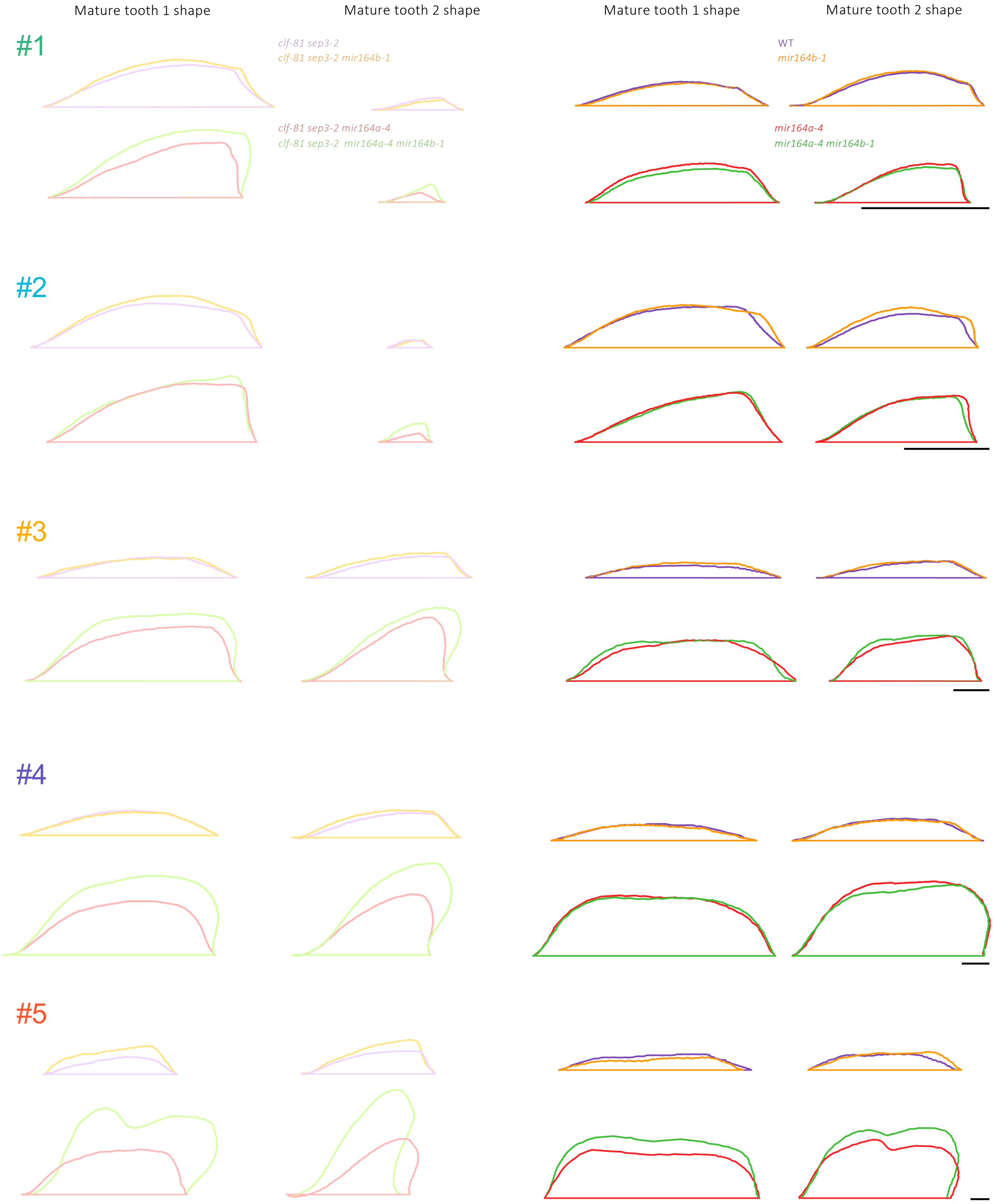


**Figure S6: Average tooth shape of mature leaves of plants under different growth conditions**
Mean shape of tooth 1 and 2 of mature leaves of the indicated genotypes under the five different growth conditions (see Figure 4b for details of conditions). All teeth of the same conditions are shown at the same scale. Bars = 1 mm

Fig S6

| Figure | Genotype | Leaf blade length (µm) | |
|---|---|---|---|
| | | @tooth 1 125µm wide | @tooth 1 250µm wide |
| 1j | WT | 483 | 885 |
| | *clf-81 sep3-2* | 498 | 917 |
| 2c | *mir164a-4* | 527 | 1074 |
| | clf-81 sep3-2 mir164a-4 | 579 | 1084 |
| 3e | *clf-81 sep3-2* | 510 | 920 |
| | *clf-81 sep3-2 mirb-1* | 530 | 977 |
| 3e | clf-81 sep3-2 mir164a-4 | 579 | 1084 |
| | *clf-81 sep3-2 mir164a-4 mir164b-1* | 580 | 1207 |
| S3a | WT | 483 | 981 |
| | *mir164b-1* | 506 | 989 |
| S3a | *mir164a-4* | 527 | 1074 |
| | *mir164a-4 mir164b-1* | 494 | 1082 |

**Table S1. Leaf size corresponding to tooth 125 or 250 µm wide**

| Figure | Growth conditions | Gentoypes | tooth 1 width (µm) | tooth 2 width (µm) |
|---|---|---|---|---|
| 1c | #3 | WT and *clf81 sep3-2* | 6600 | 5727 |
| 3h | #3 | *clf81 sep3-2* and *clf81 sep3-2 mir164b-1* | 6608 | 5641 |
| | | clf81 sep3-2 *mir164a-4* and *clf81 sep3-2 mir164a-4 mir164b-1* | 7115 | 4641 |
| S3c | #3 | *WT* and *mir164b-1* | 7134 | 6641 |
| | | *mir164a-4* and *mir164a-4 mir164b-1* | 8423 | 6678 |
| S3c | | *clf-29* and *clf-29 mir164b-1* | 4944 | 4612 |
| | | *clf-29 mir164a-4* and *clf-29 mir164a-4 mir164b-1* | 5942 | 4506 |
| S6 | #1 | *WT* and *mir164b-1* | 1807 | 1850 |
| | | *mir164a-4* and *mir164a-4 mir164b-1* | 1822 | 1460 |
| | | *clf-29* and *clf-29 mir164b-1* | 2176 | 862 |
| | | *clf-29 mir164a-4* and *clf-29 mir164a-4 mir164b-1* | 1836 | 610 |
| | #2 | *WT* and *mir164b-1* | 2712 | 2122 |
| | | *mir164a-4* and *mir164a-4 mir164b-1* | 2507 | 1951 |
| | | *clf-29* and *clf-29 mir164b-1* | 2810 | 546 |
| | | *clf-29 mir164a-4* and *clf-29 mir164a-4 mir164b-1* | 2550.000 | 644.000 |
| | #4 | *WT* and *mir164b-1* | 5548 | 4833 |
| | | *mir164a-4* and *mir164a-4 mir164b-1* | 6538 | 4344 |
| | | *clf-29* and *clf-29 mir164b-1* | 5696 | 4749 |
| | | *clf-29 mir164a-4* and *clf-29 mir164a-4 mir164b-1* | 6154 | 4285 |
| | #5 | *WT* and *mir164b-1* | 7276 | 5689 |
| | | *mir164a-4* and *mir164a-4 mir164b-1* | 8104 | 5680 |
| | | *clf-29* and *clf-29 mir164b-1* | 5029 | 5049 |
| | | *clf-29 mir164a-4* and *clf-29 mir164a-4 mir164b-1* | 6225 | 4650 |

**Table S2. Tooth width of mature leaves**

| Figure | Variable | Condition | Genotype | n | Test | p-value | Conclusion |
|---|---|---|---|---|---|---|---|
| Fig 1b | LDI | Mature | WT | 20 | Student's t-test | 8.77E-05 | *** |
| Fig 1b | LDI | Mature | *clf-81 sep3-2* | 22 | | | |
| Fig 1c | TAR | Tooth Position 1 | WT | 36 | Student's t-test | 2.36E-02 | * |
| Fig 1c | TAR | Tooth Position 1 | *clf-81 sep3-2* | 44 | | | |
| Fig 1c | TAR | Tooth Position 2 | WT | 36 | Student's t-test | 3.18E-05 | *** |
| Fig 1c | TAR | Tooth Position 2 | *clf-81 sep3-2* | 44 | | | |
| Fig 1h | pCUC2:RFP Intensity | [250-750] | WT | 18 | Student's t-test | 1.27E-03 | *** |
| Fig 1h | pCUC2:RFP Intensity | [250-750] | *clf-81 sep3-2* | 21 | | | |
| Fig 1h | pCUC2:RFP Intensity | ]750-1250] | WT | 8 | Student's t-test | 3.88E-04 | *** |
| Fig 1h | pCUC2:RFP Intensity | ]750-1250] | *clf-81 sep3-2* | 18 | | | |
| Fig 1i | CUC2-VENUS quantity | [250-750] | WT | 51 | Student's t-test | 3.47E-01 | NS |
| Fig 1i | CUC2-VENUS quantity | [250-750] | *clf-81 sep3-2* | 33 | | | |
| Fig 1i | CUC2-VENUS quantity | ]750-1250] | WT | 14 | Student's t-test | 9.10E-01 | NS |
| Fig 1i | CUC2-VENUS quantity | ]750-1250] | *clf-81 sep3-2* | 6 | | | |
| Fig 1k | LDI | [250-750] | WT | 95 | Student's t-test | 5.59E-02 | NS |
| Fig 1k | LDI | [250-750] | *clf-81 sep3-2* | 50 | | | |
| Fig 1k | LDI | ]750-1250] | WT | 47 | Student's t-test | 9.93E-01 | NS |
| Fig 1k | LDI | ]750-1250] | *clf-81 sep3-2* | 31 | | | |
| Fig 2a | pMIR164A:RFP Intensity | [250-750] | WT | 105 | Student's t-test | 1.41E-05 | *** |
| Fig 2a | pMIR164A:RFP Intensity | [250-750] | *clf-81 sep3-2* | 101 | | | |
| Fig 2a | pMIR164A:RFP Intensity | ]750-1250] | WT | 48 | Student's t-test | 9.64E-01 | NS |
| Fig 2a | pMIR164A:RFP Intensity | ]750-1250] | *clf-81 sep3-2* | 39 | | | |
| Fig 2b | CUC2-VENUS quantity | [250-750] | *mir164a-4* | 19 | Student's t-test | 3.01E-02 | * |
| Fig 2b | CUC2-VENUS quantity | [250-750] | *clf-81 sep3-2 mir164a-4* | 27 | | | |
| Fig 2b | CUC2-VENUS quantity | ]750-1250] | *mir164a-4* | 25 | Student's t-test | 9.75E-01 | NS |
| Fig 2b | CUC2-VENUS quantity | ]750-1250] | *clf-81 sep3-2 mir164a-4* | 26 | | | |
| Fig 2d | LDI | [250-750] | *mir164a-4* | 28 | Student's t-test | 1.80E-02 | * |
| Fig 2d | LDI | [250-750] | *clf-81 sep3-2 mir164a-4* | 53 | | | |
| Fig 2d | LDI | ]750-1250] | *mir164a-4* | 17 | Student's t-test | 9.22E-04 | *** |
| Fig 2d | LDI | ]750-1250] | *clf-81 sep3-2 mir164a-4* | 24 | | | |
| Fig 3f | LDI | [250-750] | *clf-81 sep3-2* | 75 | ANOVA followed by Tukey HSD | 2.26E-14 | a |
| Fig 3f | LDI | [250-750] | *clf-81 sep3-2 mir164b-1* | 38 | | | a |
| Fig 3f | LDI | [250-750] | *clf-81 sep3-2 mir164a-4* | 53 | | | b |
| Fig 3f | LDI | [250-750] | *clf-81 sep3-2 mir164a-4 mir164b-1* | 50 | | | b |
| Fig 3f | LDI | ]750-1250] | *clf-81 sep3-2* | 44 | ANOVA followed by Tukey HSD | <2e-16 | a' |
| Fig 3f | LDI | ]750-1250] | *clf-81 sep3-2 mir164b-1* | 15 | | | a' |
| Fig 3f | LDI | ]750-1250] | *clf-81 sep3-2 mir164a-4* | 24 | | | b' |
| Fig 3f | LDI | ]750-1250] | *clf-81 sep3-2 mir164a-4 mir164b-1* | 21 | | | c' |
| Fig 3g | CUC2-VENUS quantity | [250-750] | *clf-81 sep3-2 mir164a-4* | 45 | Student's t-test | 1.11E-02 | * |
| Fig 3g | CUC2-VENUS quantity | [250-750] | *clf-81 sep3-2 mir164a-4 mir164b-1* | 35 | | | |
| Fig 3g | CUC2-VENUS quantity | ]750-1250] | *clf-81 sep3-2 mir164a-4* | 23 | Student's t-test | 1.68E-04 | *** |
| Fig 3g | CUC2-VENUS quantity | ]750-1250] | *clf-81 sep3-2 mir164a-4 mir164b-1* | 34 | | | |
| Fig 3i | LDI | Mature | *clf-81 sep3-2* | 22 | ANOVA followed by Tukey HSD | <2e-16 | a |
| Fig 3i | LDI | Mature | *clf-81 sep3-2 mir164b-1* | 23 | | | a |
| Fig 3i | LDI | Mature | *clf-81 sep3-2 mir164a-4* | 23 | | | b |
| Fig 3i | LDI | Mature | *clf-81 sep3-2 mir164a-4 mir164b-1* | 23 | | | c |
| Fig S3b | LDI | [250-750] | WT | 42 | ANOVA followed by Tukey HSD | 1.25E-05 | ab |
| Fig S3b | LDI | [250-750] | *mir164b-1* | 24 | | | a |
| Fig S3b | LDI | [250-750] | *mir164a-4* | 28 | | | c |
| Fig S3b | LDI | [250-750] | *mir164a-4 mir164b-1* | 19 | | | bc |
| Fig S3b | LDI | ]750-1250] | WT | 15 | ANOVA followed by Tukey HSD | 1.39E-14 | a' |
| Fig S3b | LDI | ]750-1250] | *mir164b-1* | 12 | | | a' |
| Fig S3b | LDI | ]750-1250] | *mir164a-4* | 17 | | | b' |
| Fig S3b | LDI | ]750-1250] | *mir164a-4 mir164b-1* | 19 | | | b' |
| Fig S3d | LDI | Mature | WT | 21 | ANOVA followed by Tukey HSD | <2e-16 | a |
| Fig S3d | LDI | Mature | *mir164b-1* | 25 | | | a |
| Fig S3d | LDI | Mature | *mir164a-4* | 29 | | | b |
| Fig S3d | LDI | Mature | *mir164a-4 mir164b-1* | 27 | | | b |
| Fig S3f | LDI | Mature | *clf-29* | 26 | ANOVA followed by Tukey HSD | <2e-16 | a |
| Fig S3f | LDI | Mature | *clf-29 mir164b-1* | 31 | | | a |
| Fig S3f | LDI | Mature | *clf-29 mir164a-4* | 27 | | | b |
| Fig S3f | LDI | Mature | *clf-29 mir164a-4 mir164b-1* | 34 | | | c |
| Fig 4d | Nomalized LDI | #1 | *clf-81 sep3-2* | 19 | ANOVA followed by Tukey HSD | 1.31E-12 | a |
| Fig 4d | Nomalized LDI | #1 | *clf-81 sep3-2 mir164b-1* | 19 | | | a |
| Fig 4d | Nomalized LDI | #1 | *clf-81 sep3-2 mir164a-4* | 20 | | | b |
| Fig 4d | Nomalized LDI | #1 | *clf-81 sep3-2 mir164a-4 mir164b-1* | 20 | | | c |
| Fig 4d | Nomalized LDI | #2 | *clf-81 sep3-2* | 19 | ANOVA followed by Tukey HSD | <2e-16 | a |
| Fig 4d | Nomalized LDI | #2 | *clf-81 sep3-2 mir164b-1* | 19 | | | a |
| Fig 4d | Nomalized LDI | #2 | *clf-81 sep3-2 mir164a-4* | 18 | | | b |
| Fig 4d | Nomalized LDI | #2 | *clf-81 sep3-2 mir164a-4 mir164b-1* | 17 | | | c |
| Fig 4d | Nomalized LDI | #3 | *clf-81 sep3-2* | 11 | ANOVA followed by Tukey HSD | 1.36E-15 | a |
| Fig 4d | Nomalized LDI | #3 | *clf-81 sep3-2 mir164b-1* | 10 | | | a |
| Fig 4d | Nomalized LDI | #3 | *clf-81 sep3-2 mir164a-4* | 10 | | | b |
| Fig 4d | Nomalized LDI | #3 | *clf-81 sep3-2 mir164a-4 mir164b-1* | 9 | | | c |
| Fig 4d | Nomalized LDI | #4 | *clf-81 sep3-2* | 22 | ANOVA followed by Tukey HSD | <2e-16 | a |
| Fig 4d | Nomalized LDI | #4 | *clf-81 sep3-2 mir164b-1* | 23 | | | a |
| Fig 4d | Nomalized LDI | #4 | *clf-81 sep3-2 mir164a-4* | 23 | | | b |
| Fig 4d | Nomalized LDI | #4 | *clf-81 sep3-2 mir164a-4 mir164b-1* | 23 | | | c |
| Fig 4d | Nomalized LDI | #5 | *clf-81 sep3-2* | 5 | ANOVA followed by Tukey HSD | 3.18E-07 | a |
| Fig 4d | Nomalized LDI | #5 | *clf-81 sep3-2 mir164b-1* | 6 | | | ab |
| Fig 4d | Nomalized LDI | #5 | *clf-81 sep3-2 mir164a-4* | 5 | | | b |
| Fig 4d | Nomalized LDI | #5 | *clf-81 sep3-2 mir164a-4 mir164b-1* | 4 | | | c |
| Fig 4e | Nomalized LDI | #1 | WT | 15 | ANOVA followed by Tukey HSD | 1.08E-10 | a |
| Fig 4e | Nomalized LDI | #1 | *mir164b-1* | 15 | | | a |
| Fig 4e | Nomalized LDI | #1 | *mir164a-4* | 17 | | | b |
| Fig 4e | Nomalized LDI | #1 | *mir164a-4 mir164b-1* | 18 | | | b |
| Fig 4e | Nomalized LDI | #2 | WT | 14 | ANOVA followed by Tukey HSD | 1.05E-12 | a |
| Fig 4e | Nomalized LDI | #2 | *mir164b-1* | 13 | | | a |
| Fig 4e | Nomalized LDI | #2 | *mir164a-4* | 19 | | | b |
| Fig 4e | Nomalized LDI | #2 | *mir164a-4 mir164b-1* | 20 | | | c |
| Fig 4e | Nomalized LDI | #3 | WT | 11 | ANOVA followed by Tukey HSD | 4.78E-15 | a |
| Fig 4e | Nomalized LDI | #3 | *mir164b-1* | 10 | | | a |
| Fig 4e | Nomalized LDI | #3 | *mir164a-4* | 10 | | | b |
| Fig 4e | Nomalized LDI | #3 | *mir164a-4 mir164b-1* | 10 | | | b |
| Fig 4e | Nomalized LDI | #4 | WT | 21 | ANOVA followed by Tukey HSD | <2e-16 | a |
| Fig 4e | Nomalized LDI | #4 | *mir164b-1* | 25 | | | a |
| Fig 4e | Nomalized LDI | #4 | *mir164a-4* | 29 | | | b |
| Fig 4e | Nomalized LDI | #4 | *mir164a-4 mir164b-1* | 27 | | | b |
| Fig 4e | Nomalized LDI | #5 | WT | 5 | ANOVA followed by Tukey HSD | 8.16E-07 | a |
| Fig 4e | Nomalized LDI | #5 | *mir164b-1* | 5 | | | a |
| Fig 4e | Nomalized LDI | #5 | *mir164a-4* | 5 | | | b |
| Fig 4e | Nomalized LDI | #5 | *mir164a-4 mir164b-1* | 6 | | | b |

**Table S3. Statistical analysis**

| Locus | F primer | R primer |
|---|---|---|
| LEC2 | CCTGTTGATCCTTGCCATCT | TGAATCCTCAGCCGGTTTAC |
| UB10 | AACAATTGGAGGATGGTCGT | GTGTCGGAGCTTTCCACTTC |
| CUC2-1 (promoter) | AAACAAGAGCCCATTCCTCGT | AAAGAGAGCAAAGCCAAAGCAG |
| CUC2-2 (close to the TSS) | ATGGCGGAGACAGCCAATATC | AGAAGCAACCGTCGAGGACT |
| CUC2-3 (end of the gene) | GTAGCACCAACACAACCGTC | CTAAGCCCAAGGCCGTAGTAG |
| MIR164B-1 (promoter) | TGTACATCGGTGTGGTCTTGT | GCTGACCCATGAACATGCTA |
| MIR164B-2 (miR coding region) | CCCAGTTATGTGGTCGGAGAG | CACGCATTTGTGGTGAGAGTG |
| MIR164B-3 | GCGAGTATTAGACAGGAGATCCAA | AAGGGGCAACCAAAAGGGTTT |
| MIR164C-1 (promoter) | TCCTTCTGTCATGTTCAAAGGTC | ATTCCAACCCTCACGACGTA |
| MIR164C-2 (miR coding region) | CTTGATGGAGAAGCAGGGCAC | GAGAGACACGTGTTGGAGTAGT |
| MIR164C-3 | ACGATTTTGGCTCACGTCATTG | CCCCCGCTTTCTAAGGCAAC |

**Table S4. Primers used in this work**
